# A Multiple Regression Assessment of the Biomineral Urease Activity from Urine Drainpipes of California Rest Areas

**DOI:** 10.1101/2021.02.18.431895

**Authors:** Kahui Lim, Harold Leverenz, Cara Wademan, Samantha Barnum

## Abstract

Clogging and odor is strongly associated with ureolytic biomineralization in waterless and low-flow urinal drainage systems in high usage settings. These blockages continue to hinder widespread waterless and low-flow urinal adoption due to subsequent high maintenance requirements and hygiene concerns. Through field observations, hypothesis testing, and multiple regression analysis, this study attempts to characterize, for the first time, the ureolytic activity of the biomineralization found in alternative technologies located at 9 State-owned restrooms. Multiple regression analysis (*n* = 55, df = 4, *R*^*2*^ = 0.665) suggests that intrasystem sampling location 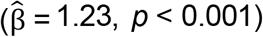, annual users per rest area 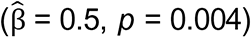, and the organic/inorganic mass fraction 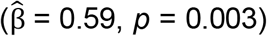, are statistically significant influencers of the ureolytic activity of biomineral samples (*p* < 0.05). Conversely, *ureC* gene abundance (*p* = 0.551), urinal type (*p* = 0.521) and sampling season (*p* = 0.956) are not significant predictors of biomineral ureolytic activity. We conclude that high concentrations of the urease alpha subunit, *ureC*, which can be interpreted as proxy measure of a strong, potentially ureolytic community, does not necessarily mean that the gene is being expressed. Future studies should assess *ureC* transcriptional activity to measure gene expression rather than gene abundance to assess the relationship between environmental conditions, their role in transcription, and urease activities. In sum, this study presents a method to characterize biomineral ureolysis and establishes baseline values for future ureolytic inhibition treatment studies that seek to improve the usability of urine collection and related source separation technologies.

## 1. Introduction

Waterless and low-flow urinals reduce water consumption, improve hygiene with touchless operation, and can be used for source separation of urine; additionally, waterless systems require less plumbing than conventional systems. However, these source-separation technologies are susceptible to biomineralization [1,2]. Biomineralization, usually of a mixed composition of struvite, calcium phosphate, calcium oxalate, and calcium carbonate, has plagued urine diversion projects since the earliest projects were studied, leading to clogging, odor, and overall user dissatisfaction [1–4].

Researchers have described the formation of biomineralization in terms of (a) cellular activities, (b) passive formation of crystals caused by biofilms, and (c) biological and chemical facilitation of crystal supersaturation conditions [5–7]. Biomineralization in urine source-separation contexts is likely governed by a combination of mechanisms.

Urease and its ureolytic activity are measures of biomineralization potential because the rate of precipitation is dependent, in part, on the rate of increase of media pH, which depends on the rate of ureolysis. The elevated pH resulting from ureolysis plays a critical role in the supersaturation crystal formation process. Because urinals are subject to intermittent supplements of a urea and an ion source, urinals and urine drainage traps become a selective breeding ground for ureolytic organisms that cause an increase in the pH of collected urine and facilitate mineral precipitation as has been observed in urological devices[8]. Ureolytic bacteria responsible for the biomineralization use the nickel-dependent metalloenzyme, urease, to catalyze the hydrolysis of urea into ammonia and bicarbonate which in turn raises the pH and creates conditions favorable of precipitation [4]. Broomfield et al. (2009), in their catheter study, demonstrated that rates of calcium and magnesium encrustation caused by various ureolytic bacteria isolates is correlated with an increase in ureolytic activity [9] An elevated pH promotes calcium phosphate and oxalate stone formation due to a shift in phosphate speciation from HPO_4_^2-^to PO_4_^3-^and the decomposition of ascorbic acid into oxalate—both cases represent an increase in ion concentrations that lead to elevated encrustation rates found in catheters [10]. Ureolysis also results in carbonate and bicarbonate ion formation which can further contribute to biomineralization as the urine becomes supersaturated [11]. Researchers similarly showed that greater ureolytic rates from bacterial urease are correlated with greater rates of calcium carbonate precipitation [12–14]. Studies using *Proteus mirabilis* have shown that urease defective mutants fail to form crystalline biofilms in laboratory models, demonstrating the key role of pH and urease activity in crystal formation [15]. In dental plaque studies, researchers suggest that ammonia generating capacity in a mixed-species model of ureolytic oral biofilms is essential for the stabilization of microbial communities in ureolytic environments [16]. Losses of sufficient quantities of urease resulted in the acidification of biofilms and a decrease in community diversity [16].

Through multiple linear regression modelling, this study will be the first of its kind to: (a) model biomineral enzyme activity in terms of both categorical and quantitative predictors, (b) examine biomineral enzyme activity from urine source-separation technology, and (c) do so on a geographic scale with a sufficiently large sample size. This study also builds upon previous works describing soil or biofilm ureolytic activity that (a) use small sample sizes in parametric hypothesis tests (n=6) or multiple regression (n=4), (b) neglect discussion of model validation beyond the coefficient of determination (R^2^), (c) do not discuss whether their data fits assumptions required for application of a statistical test, and (d) mention statistical significance, but not practical significance, i.e. the magnitude of effect [17–20].

Finding a link between environmental parameters such as intrasystem sampling location, usage frequency, seasonality, gene abundance found through qPCR, and urinal types with the enzymatic activity of the biomineral samples will be useful in understanding the effects of restroom configuration on ureolytic activity. Understanding the effect of seasonality and sampling locations within a urine drainage system where ureolytic activity is highest may be insightful when predicting locations and times of year where the components of the urine collection system are most susceptible to biomineral fouling.

## 2. Materials and Methods

The coming subsections will describe the sampling procedures and locations followed by methods used in downstream analyses to quantify the environmental variables used in the statistical analyses. The R Markdown HTML output containing the script can be found in the Online Resources section. The raw environmental data can be found in the Dryad repository (DOI:10.25338/B82906) as an .RDS file.

### 2.1 Sample Collection

Rest areas were categorized by the types of urinals installed: conventional ∼ 1 gal/flush, low-flow ∼0.125 gal/flush, and waterless no flush. Biomineralization deposits were scraped into sterile 50 mL conical tubes from fouled fixtures and drainage systems when available. A total of 2 conventional, 2 low-flow, and 5 waterless public restrooms along California highways, also known as rest areas, were observed in this study. A summary of sites, drainpipe configurations, and characteristic samples are shown in Figure 1 as: (a) location of sampling sites with respect to urinal type used in this study, (b) biomineralization formation on a waterless urinal cartridge at Erreca on 16 Sep 2019, (c) a view of reduced internal pipe diameter by biomineralization in a urine drainage pipe at the Dunnigan northbound oriented public rest area on 12 Dec 2019, and (d) general drainage system layout consisting of the drains directly connected to the urinals, which collectively flows into a main drain also connected to the sink drains. The men’s restrooms were typically fitted with two urinals at two different heights to conform to the American Disability Act (ADA).

**Fig. 1.**
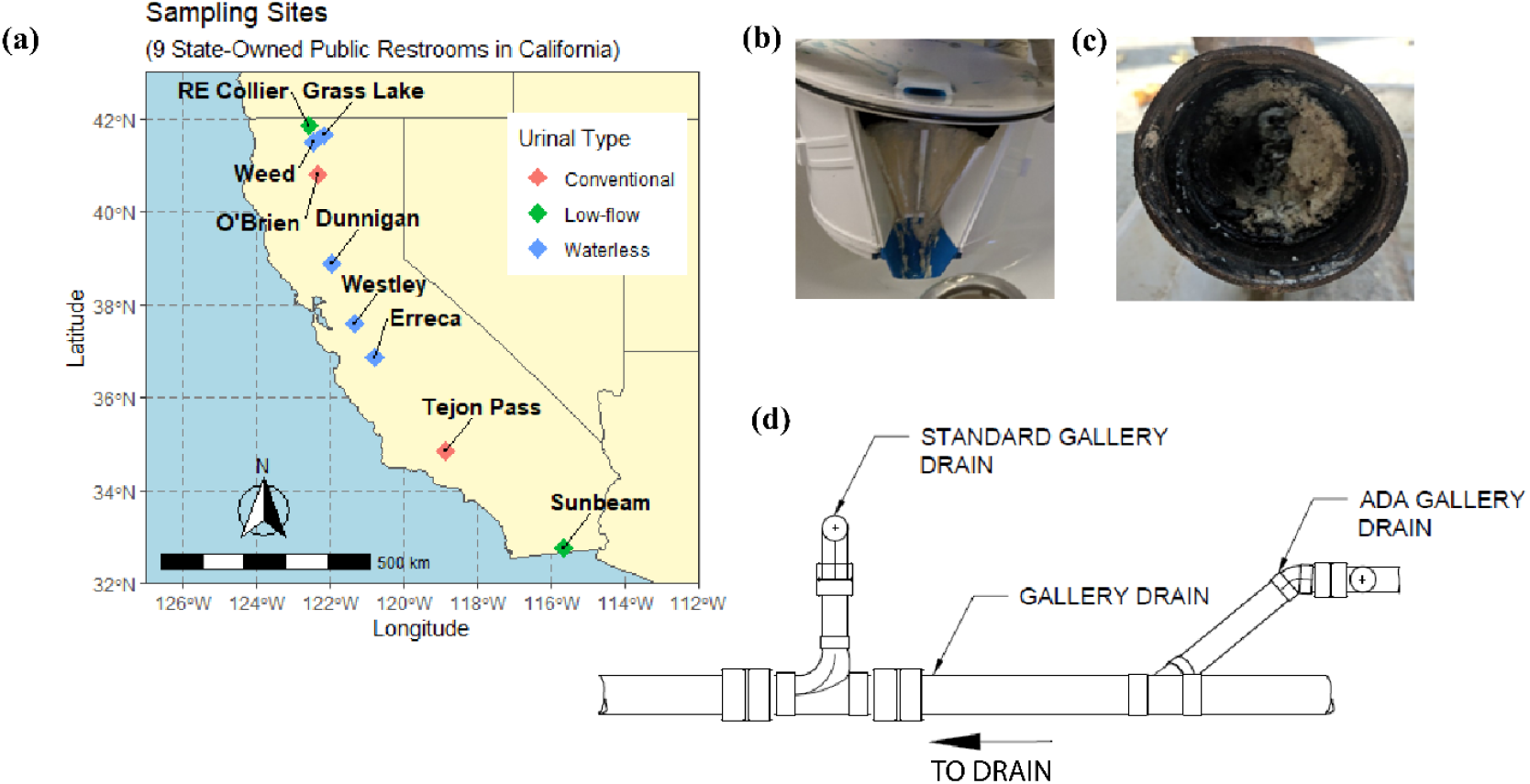
Sampling sites, characteristic samples, and typical drainage system configurations

All samples were stored in an ice chest after collection and processed within three days of sample collection. Previous work monitoring the ureolysis rate in soils have found that a distinct slowdown in ureolytic rate was not detected until 8 months of cooled storage [21]. As such, the sampling preservation measures were deemed adequate.

### 2.2 Biomineral Ureolytic Enzyme Activity Characterization

To compare enzymatic activities of biomineral samples between various sites *in vitro*, a known wet mass of the biomineral samples was suspended and mixed in a 100 mL volume of 7.3 pH 200 mM (4-(2-hydroxyethyl)-1-piperazineethanesulfonic acid) (HEPES) buffer containing 2.5% urea m/m. The rate of increase in conductivity is proportional to that of urea hydrolysis and can be used as a surrogate measure for enzymatic activity [22]. As a comparative basis between samples, one unit of specific activity is defined as uS cm^-1^min^-1^ per gram of volatile solids (VS).

Gravimetric analyses followed standard methods for the examination of water and wastewater [23]. A mass balance was performed by comparing the wet solid mass with the dry mass following 105°C desiccation and fixed mass after 550°C ashing. Total solids (TS) represent the inorganic matter in the sample while VS represents organic matter. Each biomineral sample was analyzed in triplicate and then averaged.

### 2.3 Quantifying Gene Abundance using Real-time Polymerase Chain Reaction (qPCR)

To examine the relationship between *in vitro* ureolytic activity and the genetic predispositions for ureolysis, the genomes of phylotype representatives for the presence of urease genes were examined by qPCR. A similar protocol was described previously [24]. The urease associated gene were designed on the urease alpha subunit encoding gene (*ureC*). Primer sequences were obtained from the literature [25]. Sensitivity and efficiency were established from the y-intercept and slope of the standard curve, which was created by running triplicate, 10-fold serial dilutions of plasmid DNA containing the ligated amplicon of each gene (Eurofins Genomics LLC, Louisville, KY). The sensitivity of ureC-F (TGGGCCTTAAAATHCAYGARGAYTGGG) and ureC-R

(SGGTGGTGGCACACCATNANCATRTC) was <4,000 copies/qPCR reaction and the efficiency was 80.6% (R^2^ = 0.9974). Poor sensitivity and low efficiency for *ureC* is expected due to the nature of SYBR degenerative primers. Biomineral samples were kept frozen at −20°C prior to DNA extraction. DNA was manually extracted from 0.25 g of sample using a commercially available kit following manufacturer recommendations and eluted in 100 μL of diethylpyrocarbonate (DEPC) treated water (Qiagen DNeasy Power Soil Kit, cat # 12888-50). Each 12 μL reaction contained 6 μL SYBR master mix (Applied Biosystems SYBR Green PCR Master Mix, cat # 4309155), 0.48 μL of a primer-water mixture (primers at final concentration of 400 nM), 4.52 μL of DEPC-treated water, and 1 μL of extracted DNA. qPCR was performed using an automated fluorometer (ABI PRISM 7900 HTA FAST, Thermo Fisher Scientific). Standard amplification conditions were used: 95°C for 3 min, 40 cycles of 95°C for 15 s, 52°C for 30 s, and 72°C for 30 s, with a melting curve at 95°C for 15 s, 52°C for 15 s, and 95°C for 15 s. Data was analyzed using Applied Biosystems SDS software, version 2.4. Fluorescent signals were collected during the annealing phase and C_q_ values extracted with a threshold of 0.2 and baseline values of 3–10 for the *ureC* assay. Amplification specificity was verified using the dissociation temperature (T_m_) of the qPCR amplicons specific to each gene. Acceptable T_m_ ranges were determined to be +/- 2% of the positive controls. For *ureC*, the acceptable T_m_ range was 80.8°C - 84.1°C. Samples with detectable amplification but with T_m_’s outside of the acceptable ranges were considered false positives and were deemed negative for the gene of interest. The absolute copy numbers were also normalized in terms of volatile solids (VS) mass present in the biomineral samples.

### 2.4 Statistical Analyses

All statistical work and data visualization was done using R version 4.0.2. An a-priori power analysis was first used to inform the design of this study, suggesting that a linear model can sufficiently capture a large effect size (f=0.35) at a level of significance of 0.05 for a power of 0.8 using 1 tested dependent variable and 5 total predictors with a minimum sample size of 25 [26]. After excluding sample rows missing data from low quality qPCR reads and samples that did not have enough mass for gravimetric analysis or biomineral enzyme activity, this randomly sampled, complete case analysis included a sample size of 55 from 9 different facilities. In the regression analysis, conventional urinals were aggregated with low-flow urinals because both urinal types include flush water. A stepwise forward variable selection method was used. A corrected Akaike information criterion (AICC) was also used to validate model selection [27]. The ordinary least squares (OLS) multiple regression analysis was performed assuming that a natural log-log transformed linear model is an adequate descriptor of the system, whereby normality was verified in the Supplementary Information section. A natural log-log transformed dataset enables for a practical interpretation of the effect size as a percent change, or in this case, the elasticity between two biological variables [28]. Regression coefficients were interpreted as natural log-level for categorical variables. For the coming subsections, unless specified, variables will be discussed in terms of natural logarithms.

### 2.5 Characterizing the Ureolytic Activity in Urinal Traps *in situ*

*In situ* urinal trap testing was conducted to characterize the ureolytic rate at the time of sample collection within the urine drain trap. *In situ* biomineral ureolytic activity was used to support the regression analysis derived from *in vitro* urease assays. The project team developed a method using pH and conductivity meters to characterize the baseline ureolytic rates. A description of the trap testing is shown graphically in Figure 2.

**Fig. 2.**
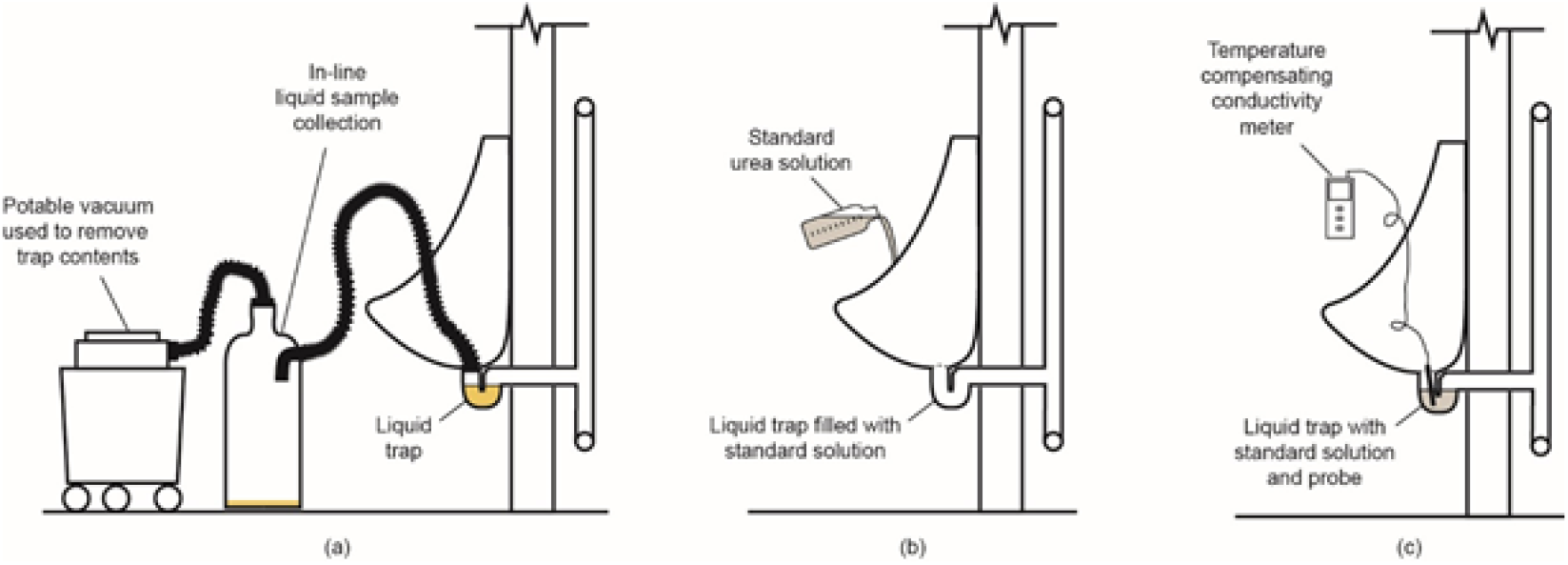
Schematic of *in situ* trap activity test procedure: (a) using portable vacuum and in-line liquid sample collector to remove trap contents, (b) application of standard urea solution to empty trap, and (c) testing of urinal liquid trap to determine relative activity

The in-situ urinal trap procedure was conducted as follows:

First, the urine drain trap is vacuumed out as shown in Figure 2. Once emptied, a 200 mM 7.3 pH HEPES buffer containing 2.5% m/m urea is added until the drain trap is full. Logging pH and EC meters were submerged in the trap opening and recorded for a total of 10 minutes from which the ureolytic rate could be estimated using the rate of EC formation.

## 3. Results and Discussion

After evaluating and selecting the most parsimonious multiple linear regression model composed of categorical and quantitative environmental variables, the observed influence, or lack thereof, of these variables will be discussed in the context of biomineral ureolytic activity.

### 3.1 Multiple Linear Model and Validation

The multiple linear model composed of 55 observations is described in Tables 1 and 2. As shown in correlation heatmaps and residual analysis from Supplementary Figures 1 and 2, the linear model is in agreement with the Gauss-Markov OLS regression assumptions, which require that: a) the expected value of the regression residuals tends towards zero, b) the residuals are homoscedastic c) there is no autocorrelation between the regressors and the residuals such that exogeneity is upheld, d) the predictors are not multicollinear, and e) the residuals are also normal [28]. The residuals shown in Supplementary Figure 1 do not appear to have a trend based on the index plot, do not exhibit any correlation with each other from the autocorrelation plot, and appear homoscedastic from the fitted values vs. residuals plot. Finally, the residuals also appear normally distributed from the quantile-quantile plot in Supplementary Figure 1. It was concluded that the natural log-log linear model appropriately describes natural logarithmically transformed data and that the model fits well with the data. The AICC model selection results are shown in Supplementary Table 2, suggesting that the most parsimonious and probable model is Model 3 [27,29].

**Table 1.** Summary of effect sizes of significant predictors on biomineral ureolytic activity

| Significant Predictor Variables | $\hat{\beta}$ | CI (95%) | Effect on Biomineral Activity per g VS as Elasticity<sup>a</sup> |
| --- | --- | --- | --- |
| Annual Users per Rest Area | 0.5 | 0.17, 0.82 | A 25% increase in annual users per rest area corresponds to a 11.7 (3.9, 20.1) % increase in biomineral activity |
| VS/TS (g/g) | 0.59 | 0.21, 0.97 | A 25% increase in VS/TS (g/g) corresponds to a 14.1 (4.8, 24.2) % increase in biomineral activity |
| Intrasystem Location: Main Drain | 1.24 | 0.83, 1.64 | Compared to samples obtained from cartridges, those obtained from the gallery main drain had a 245 (129, 416) % larger geometric mean in biomineral activity |

<sup>a</sup> Parenthetical contents represent effect sizes at limits of confidence intervals

**Table 2.** Multiple regression summary of model predicting biomineral ureolytic activity

|  | **Model 1** | **Model 2** | **Model 3** | **Model 4** | **Model 5** | **Model 6** |
| --- | --- | --- | --- | --- | --- | --- |
| Predictor Variables | Estimates | Estimates | Estimates | Estimates | Estimates | Estimates |
| Intercept | **-7.35**<br>**(-12.80 – -1.90)\*\*** | -2.18<br>(-6.92 – 2.57) | -0.55<br>(-5.05 – 3.94) | -0.46<br>(-5.00 – 4.08) | -0.56<br>(-5.12 – 3.99) | -0.36<br>(-4.92 – 4.20) |
| Annual Users per Rest Area | **0.96**<br>**(0.56 – 1.37)\*\*\*** | **0.57**<br>**(0.22 – 0.92)\*\*** | **0.50**<br>**(0.17 – 0.82)\*\*** | **0.47**<br>**(0.14 – 0.81)\*\*** | **0.49**<br>**(0.16 – 0.83)\*\*** | **0.49**<br>**(0.16 – 0.82)\*\*** |
| Intrasystem Location: Gallery Drain |  | -0.28<br>(-0.62 – 0.07) | -0.19<br>(-0.52 – 0.13) | -0.15<br>(-0.51 – 0.21) | -0.19<br>(-0.53 – 0.15) | -0.14<br>(-0.50 – 0.22) |
| Intrasystem Location: Gallery Main Drain |  | **1.02**<br>**(0.61 – 1.43)\*\*\*** | **1.24**<br>**(0.83 – 1.64)\*\*\*** | **1.26**<br>**(0.85 – 1.68)\*\*\*** | **1.24**<br>**(0.83 – 1.65)\*\*\*** | **1.23**<br>**(0.82 – 1.64)\*\*\*** |
| VS/TS (g/g) |  |  | **0.59**<br>**(0.21 – 0.97)\*\*** | **0.57**<br>**(0.19 – 0.96)\*\*** | **0.58**<br>**(0.17 – 0.99)\*\*** | **0.56**<br>**(0.18 – 0.95)\*\*** |
| ureC Concentration (copy #/g VS) |  |  |  | 0.01<br>(-0.02 – 0.04) |  |  |
| Sampling Season |  |  |  |  | 0.01<br>(-0.31 – 0.33) |  |
| Urinal Type |  |  |  |  |  | -0.12<br>(-0.49 – 0.25) |
| Observations | 55 | 55 | 55 | 55 | 55 | 55 |
| R<sup>2</sup> / R<sup>2</sup> adjusted | 0.299 / 0.286 | 0.595 / 0.571 | 0.662 / 0.635 | 0.665 / 0.630 | 0.662 / 0.628 | 0.665 / 0.631 |

<sup>a</sup> Significance codes: 0 '\*\*\*' 0.001 '\*\*' 0.05 '\*' 0.1 '.' 1

The regression results describing the most probable model (Model 3) is shown in Table 1 and 2, which also depicts the regression results from other tested models. The results presented in Tables 1 and 2 suggest that *ureC* gene concentrations (Model 4, *p* = 0.551), sampling season (Model 5, *p* = 0.956), and urinal types were statistically insignificant predictors of ureolytic activity (*p* > 0.05) and of low practical significance as indicated by the relatively small regression coefficients (see Table 6). From Table 1, the strongest predictor of biomineral ureolytic activity was the sampling location, namely, those sampled from the main urinal drainage pipes exhibited the greatest enzymatic activity. In Model 3, the second strongest predictor was the organic to inorganic fraction. Annual number of users at a given rest area also positively influenced urease activity likely due to the increased loading and usage frequency resulting in a semi-constant stream of nutrients and salts necessary for a strong ureolytic community to develop and thrive.

### 3.2 The Influence of Organic Matter on Ureolytic Activity

That the organic content is significant (*p* = 0.003) and of sizeable effect 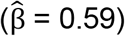 in predicting ureolytic activity, as shown in Table 1, is consistent with past findings from soil research that found correlations between organic matter concentrations and urease activity [13,14,30]. Others also observed that increased carbohydrate availability at neutral pH was correlated with increased *Actinomyces naeslundii* biofilm urease activity [14,17]. Liu et al. (2008), however, noticed that carbohydrate availability had no effect on *ureC* gene expression marked by through reverse-transcriptase quantitative real-time PCR (RT-qPCR) mRNA transcripts. Liu et al. (2008) hypothesizes that these observations were due to carbohydrate availability and pH modulation affecting the expression of genes other than *ureC* responsible for urease synthesis or apoenzyme activation [17].

Increasing the biomass of the inoculum by providing a carbon source in microbial induced calcite precipitation studies has been reported to promote the ureolytic activity [14]. Tobler et al. (2011) concluded that molasses supplementation selected for a larger microbial community that obtains their nitrogen from ureolysis, though there is no nitrogen limitation in urinals [14]. Others, who studied the environmental factors affecting microbially induced calcium precipitation concluded that increasing biomass may also increase ureolytic activity as there could be more active cells present [31]. Extracellular urease has also been suggested to be stabilized by adsorption to soil colloids, particularly organic matter, which may be similar to that observed in biomineral samples obtained from urine drain pipes [19].

One limitation of this study is that it is unclear what component of the organic fraction is correlated with increased ureolytic activity as VS is a bulk measurement encompassing any organic mass. Within the biomineral/stone matrix is also an organic fraction composed of carbohydrates, proteins, lipids, and dead cell mass that binds the mineral fraction of the precipitate [4]. Therefore, future research could evaluate different organic components such as proteins and exopolysaccharide substances (EPS).

### 3.3 The Non-effect of Urinal Type and Seasonality on Ureolytic Activity

In addition to the linear regression results, Kruskal-Wallis testing for biomineral ureolytic activity between waterless and low-flow urinals provides evidence that waterless and low-flow are likely identical in population in terms of biomineral activity (*p* = 0.47). While urinal type is not a statistically significant predictor of ureolytic activity, biomineral samples from waterless urinals have exhibited a greater maximum ureolytic activity than any biomineral sample obtained from low-flow urinals in this study, as shown on Figure 3.

**Fig. 3.**
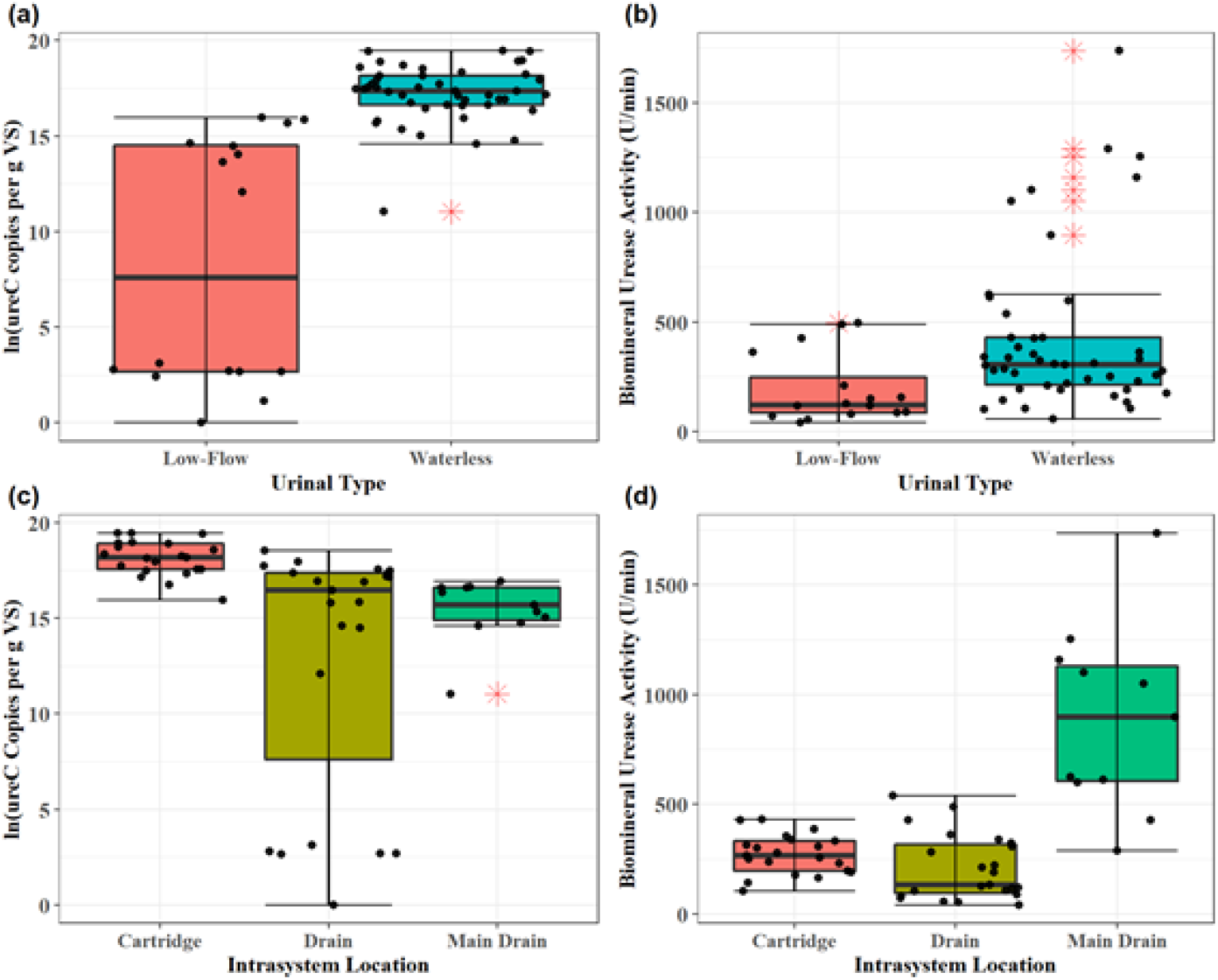
Descriptive statistics on the effects of urinal type on natural log-transformed *ureC* gene copies and biomineral activity

Finally, sampling season (as shown in Table 2) demonstrated no statistical (*p* < 0.001) or practical significance 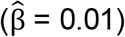 in predicting biomineral activity. This may explain why fouling is a year-round phenomenon, as the biomineral ureolytic activity remains unaffected by seasonality, as the high urease activities year-round facilitate conditions necessary for precipitation to occur. Because seasonality does not seem to impact biomineral activity, future observations on the ureolytic activity of urine drainpipes may be performed without temporal confounding effects. Though, future microbial ecology studies are needed to understand the bacterial community structure of the biomineral samples and should include sampling events from different seasons.

### 3.4 Effects of Intrasystem Sampling Location on Ureolytic Activity

While the ureolytic activity of biomineral samples obtained from the drainage pipes immediately following the drain traps were not significantly different from those corresponding to samples obtained from waterless urinal cartridges (Pairwise Wilcoxon Rank Sum: *p* = 0.053), samples taken from the main drain lines which contacts handwashing water were significantly non-identical in terms of ureolytic activity (Kruskal-Wallis: *P* < 0.001; Pairwise Wilcoxon Rank Sum: *p* < 0.001). Within one system, cartridges and gallery drain lines immediately succeeding the urinal are exposed to the same urine feed without mixing with potable water and thus face similar environmental conditions that influence ureolytic activity [13]. Because drain line samples directly follow cartridge samples and are exposed to the same urine, the relative similarity in environmental conditions between cartridge and drain line samples may explain their different ureolytic rates compared to main drainpipe samples but not with each other.

### 3.5 Biomineral Ureolytic Activity may be Predicted by Transcriptional Activity more than by *ureC* gene abundance

Kruskal-Wallis testing results suggest that the *ureC* abundance between low-flow and waterless urinals are significantly nonidentical (*p* < 0.001), but there was no detected significant effect on biomineral ureolytic activity as suggested by the multiple regression results shown in Table 2. The lack of statistical significance describing the relationship between *ureC* gene copies and ureolytic activities disagrees with bivariate correlation studies done by Fisher et al. (2016) and Sun et al. (2019), where it was found that soil ureolysis rates were significantly correlated with *ureC* gene copies. Notably, neither studies discussed effect size and used a small sample size (n < 12) for analyses describing individual soil horizons [32]. Conversely, other soil urease studies have also found that ureolytic activities are correlated with total nitrogen (TN), total carbon (TC), and soil organic carbon (SOC) concentrations, but not the abundance of *ureC* genes as in agreement with our study [33]. The regression results suggest that ureolytic gene abundance is insufficient in predicting ureolytic activity in a linear model.

Greater abundances of potentially ureolytic bacteria indicated by proxy of sample *ureC* gene concentrations, may not be correlated with biomineral ureolytic rates as suggested by the regression results. That *ureC* was detectable indicates that part of the bacterial community in the biomineral samples has the urease-positive genotype, but not all bacteria with the *ureC* may be displaying a urease-positive phenotype [34]. This is because urease activity may not be expressed under the growth conditions found in urine drain pipes, and may explain why urease activities did not differ significantly when grouped by urinal type [34]. Expression of the urease-positive genotype and the eventual translation into the urease protein is regulated at the transcriptional level rather than at the genomic level [35–37].

That *ureC* gene abundance is not a statistically significant predictor of biomineral ureolytic activity is likely due to the need for environmental conditions that would induce certain microbial transcriptional responses that cause an increase in urease activity. When comparing *ureC* copies per g VS, values grouped by intrasystem sampling location differed significantly between cartridge vs. gallery drain (Kruskal-Wallis: *p* < 0.001; Wilcoxon Rank Sum: *p* < 0.001) and cartridge vs. gallery main drain (Kruskal-Wallis: *P* < 0.001; Wilcoxon Rank Sum: *p* < 0.001). However, Figure 3 reinforces hypothesis testing results in that samples from the main drain with the lowest functional gene concentrations exhibited maximal ureolytic activity of all samples as predicted by the multiple regression model. One possible explanation is that the main drains and low-flow urinal drain lines are exposed to flush and sink water, which leads to a decrease in nitrogen concentrations in the stream contacting the biofilm due to dilution. In response, the ureolytic ammonia oxidizing bacterial community may be upregulating *ureC* transcription to produce more urease to convert the urea into ammonia at a faster rate for pH regulation or to acquire ammonia for biomass production or energy generation [38]. This hypothesis is in agreement with the literature, as researchers have shown that *ureC* mRNA transcripts were 10-fold higher in *Ruminococcus albus* cultures grown on peptides than those grown using an ammonia or urea-based media [39].

From Figure 4, the observation that conventional and low-flow urinals can have similar *in situ* ureolytic rates with those from waterless urinals is consistent with the regression results where it was found that urinal type is neither a significant (*p* = 0.521) and practical 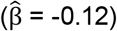 predictor of the *in vitro* biomineral ureolytic activity. Figure 4 demonstrates that Dunnigan northbound, a waterless urinal site, exhibited the greatest maximum *in situ* ureolytic rates. Conversely, Tejon Pass, an SRRA fitted with conventional urinals, ranked 2^nd^ of all sites screened for *in situ* ureolytic rate.

**Figure 4.**
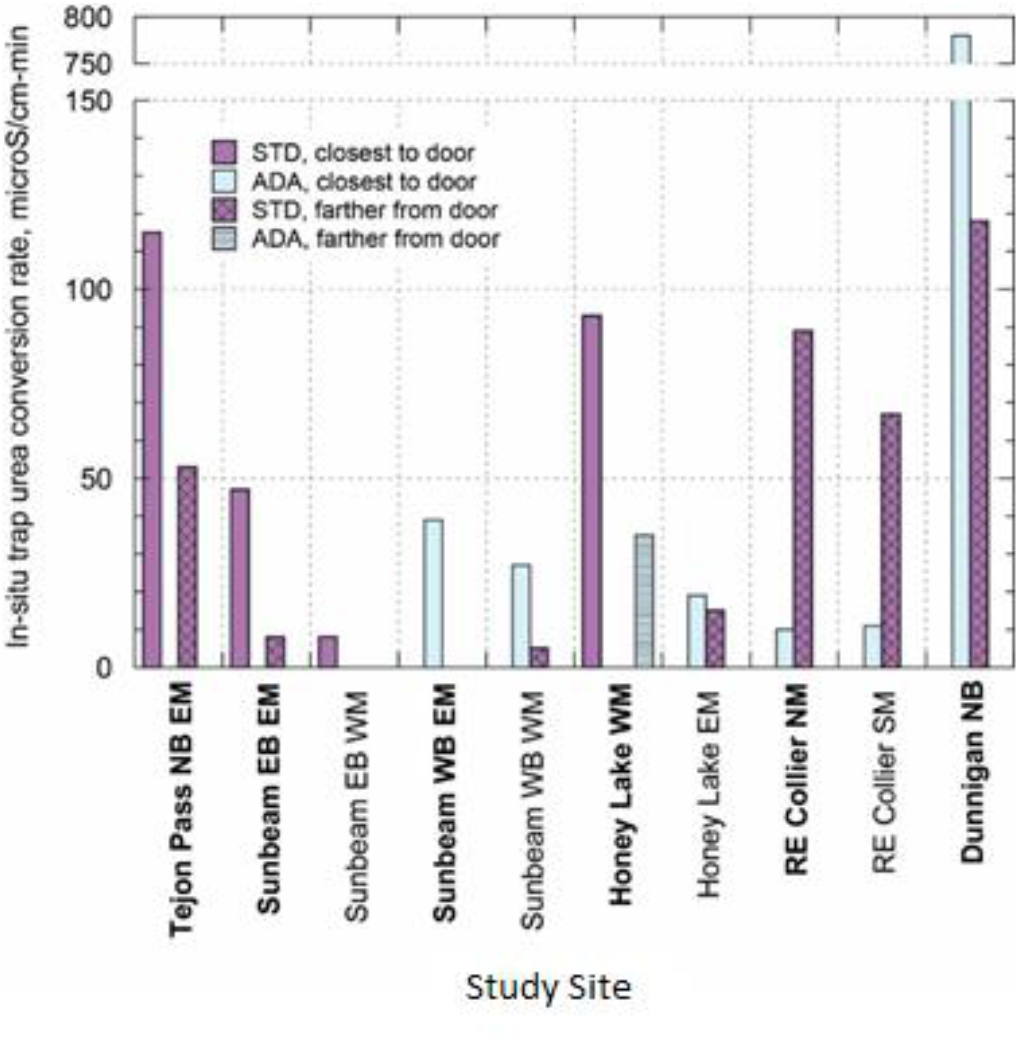
Comparison of *in situ* trap urea conversion rate for various SRRA with trap-type urinals

Our findings indicate that flush water alone may not be an adequate preventative measure for preventing ureolytic biomineralization, as urease activity can be as strong in conventional and low-flow biomineralization as it is in waterless biomineralization, even if there is a smaller ureolytic community in flush type urinals as indicated by low relative *ureC* gene concentrations shown in Figure 3. It is also possible that flush water may also influence the precipitation chemistry in drain lines, as flush water containing elevated magnesium and calcium concentrations may contribute to crystallization. While the smaller abundance of *ureC* gene concentrations in low-flow urinal samples is insufficient in accounting for the similar ureolytic activities exhibited by the two urinal types and intrasystem sampling locations, the differences in *ureC* gene concentrations grouped by urinal type shown in Figure 3 may likely be due to a difference in community structures. Future next-generation-sequencing and microbial ecology studies should visualize the potentially ureolytic microbial community structure by sequencing the *ureC* gene in addition to 16S rRNA to visualize the total bacterial community to find relationships between the bacterial community, environmental factors, and ureolytic activity.

In conjunction with measuring bulk parameters such as pH, future studies should incorporate RT-qPCR to determine the effects of nutrient concentrations on urease gene expression at the transcriptional level. A future RT-qPCR experiment on ureolytic biomineral samples can reveal how the effects of varying dilution rates between low-flow and waterless urinals affects the transcriptional activity of a gene of interest and its relationship with ureolytic activity.

## Conflicts of Interest

The authors declare no competing financial interest.

## Acknowledgements

This research was funded by the California Department of Transportation under Agreement Number 65A0734. The authors also thank the American Water Works Association CA-NV Section for their Graduate Fellowship awarded to Kahui Lim. Finally, I would like to personally thank my friend, Konrad Franco, sociologist and statistician, for his insight in developing the statistical model.

## Notes

DOI:10.25338/B82906

### Competing Interest Statement

The authors have declared no competing interest.

### Summary of Updates

Fixed special characters on abstract for bioRxiv website display.

