## Supplementary Information for "A Multiple Regression Assessment of the Biomineral Urease Activity from Urine Drainpipes of California Rest Areas"

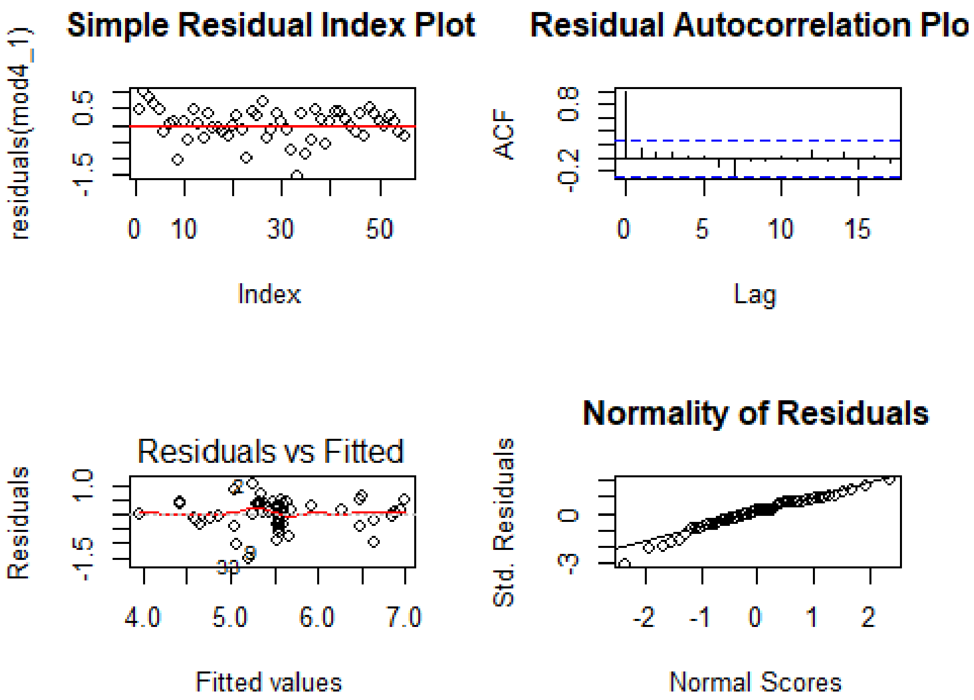


**Supplemental Figure 1** The residuals of the hypothesized model including *ureC* gene concentration as a predictor (model 4) demonstrates adherence to the Gauss-Markov assumptions of the linear model described in the manuscript.

a)

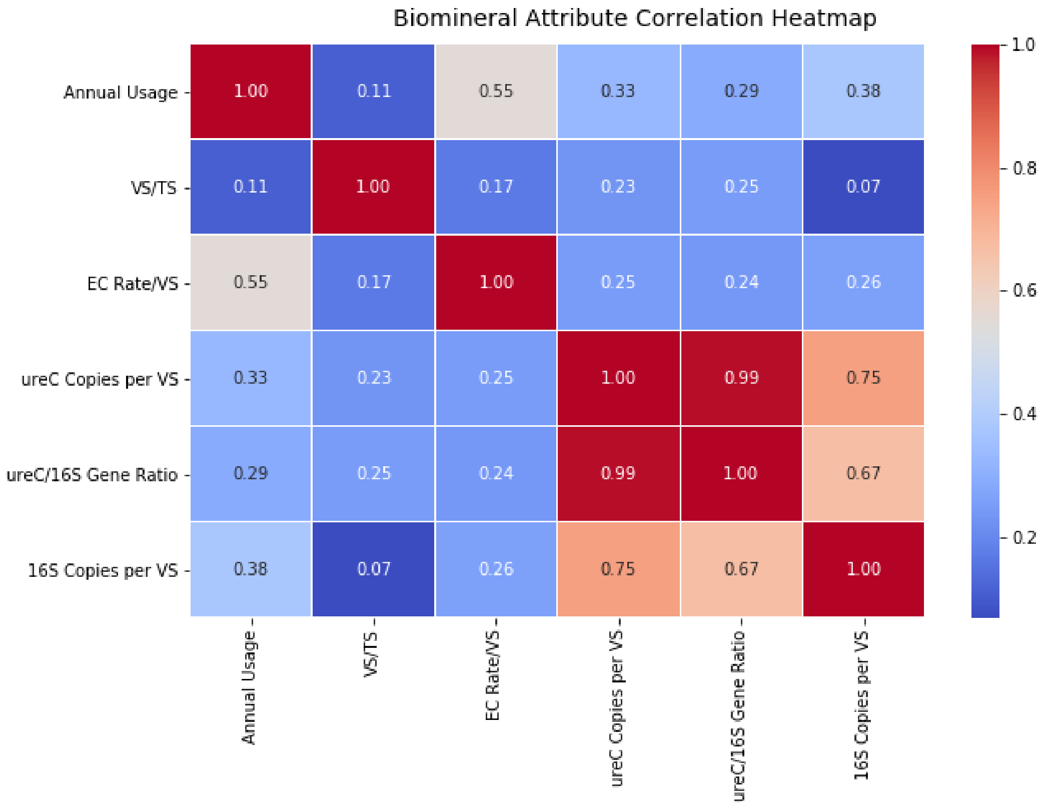


b)

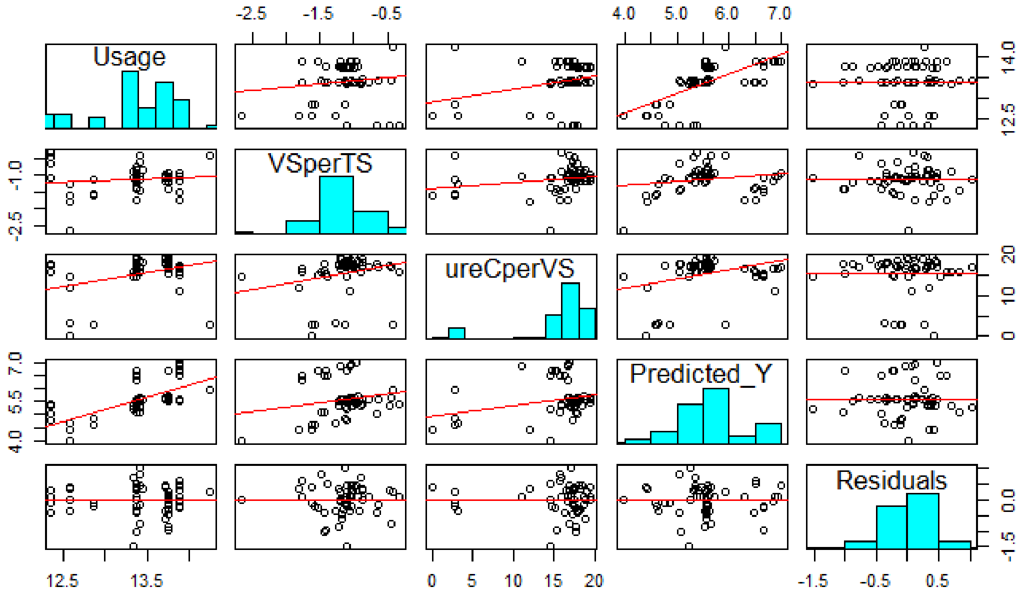


**Supplemental Figure 2a/2b** The correlation matrices suggest that the independent variables included in the multiple regression analysis is not affected by multicollinearity and that confounding factors are not observed.

**Supplemental Table 1** Summary of *in situ* trap test data

| Site | Description[a] | Date | Trap volume (mL) | *In situ* activity (uS/cm-min) | Temperature (°C) |
|---|---|---|---|---|---|
| Tejon Pass, NB | EM STD[b,c] | 9/18/2019 | 510 | 115 | 19 |
| | EM STD[b] | | 510 | 53 | 19 |
| Sunbeam EB | EM STD[b,c] | 9/17/2019 | 175 | 47 | 33 |
| | EM STD[b] | | 175 | 8 | 33 |
| | WM STD[c] | | 175 | 8 | 33 |
| Sunbeam, WB | EM ADA[b,c] | 9/17/2019 | 175 | 39 | 33 |
| | EM STD[b] | | 175 | 0 | 33 |
| | WM STD | | 175 | 5 | 33 |
| | WM ADA[c] | | 175 | 27 | 33 |
| Honey Lake | WM STD[b,c] | 12/12/2019 | 175 | 93 | 15 |
| | WM ADA[b] | | 175 | 35 | 15 |
| | EM STD | | 120 | 15 | 15 |
| | EM ADA[c] | | 120 | 19 | 15 |
| RE Collier | NM ADA[b,c] | 12/13/2019 | 130 | 10 | 12 |
| | NM STD[b] | | 200 | 89 | 12 |
| | SM ADA[c] | | 130 | 11 | 12 |
| | SM STD | | 150 | 67 | 12 |
| Dunnigan, NB | STD[b] | 12/12/2019 | 400 | 118 | 25 |
| | ADA[b,c] | | 400 | 780 | 25 |

[a] EM = eastern men's restroom, WM = western men's restroom, NM = northern men's restroom, SM = southern men's restroom, STD = standard urinal fixture height, ADA = lower urinal height to comply with American's with Disabilities Act

[b] Primary restroom facility at SRRA

[c] Urinal located nearest to door

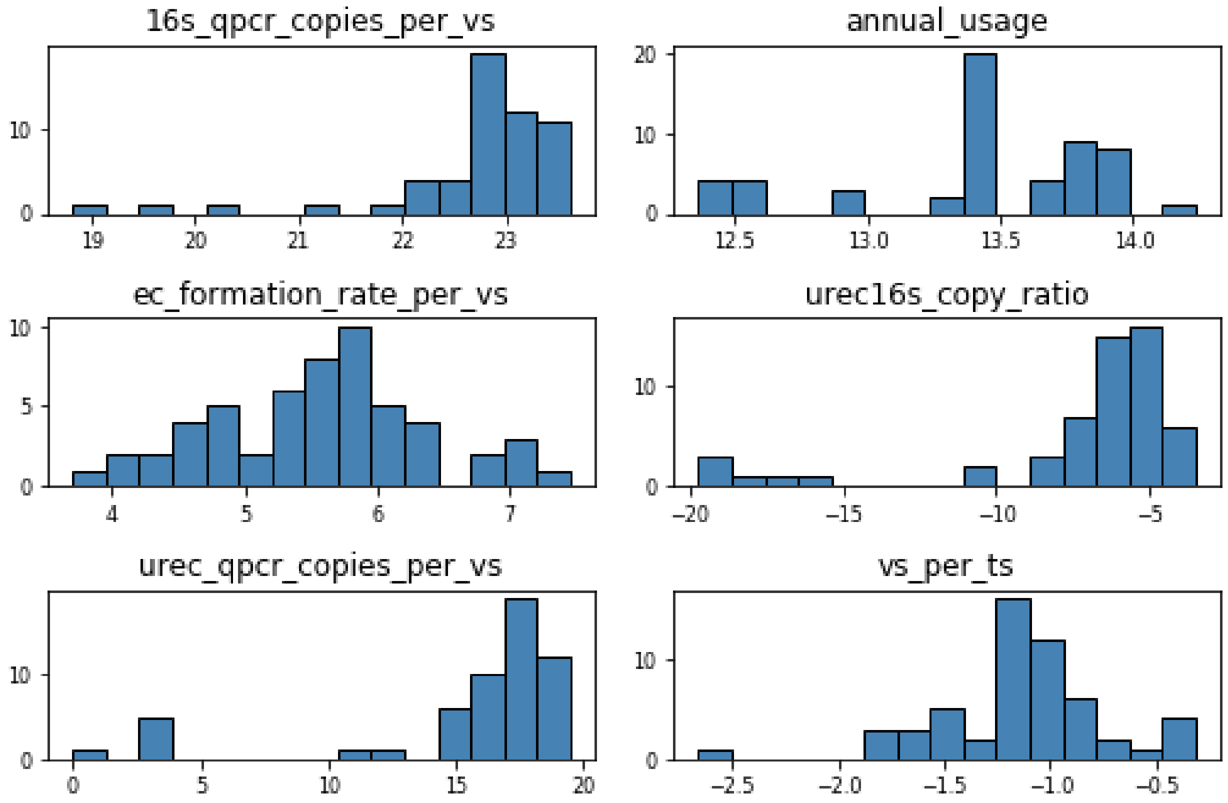


**Supplemental Figure 3:** Histograms depicting the data distribution of natural logarithmically transformed data set.

**Supplemental Table 2** Akaike Information Criterion Results for Model Selection

| **Model** | **Delta** | **df** | **Weight** |
|---|---|---|---|
| Model 3 | 0 | 6 | 0.5097 |
| Model 6 | 2.148 | 7 | 0.1741 |
| Model 4 | 2.233 | 7 | 0.1669 |
| Model 5 | 2.628 | 7 | 0.137 |
| Model 2 | 7.437 | 5 | 0.01237 |
| Model 1 | 32.85 | 3 | 3.751e-08 |
